# INSM1 Regulates Neuroendocrine Plasticity and Tumor Progression in Prostate Cancer

**DOI:** 10.64898/2026.07.30.741780

**Authors:** Chiachen Chen, Siyuan Cheng, Lin Li, Jeyaluxmy Sivalingam, Xin Gu, Yunshin Yeh, Xiuping Yu, Michael S. Lan

**Affiliations:** Department of Genetics, Louisiana State University Health Sciences Center, New Orleans, LA 70112, USA; Department of Molecular Biology and Biochemistry, Louisiana State University Health, Shreveport LA 71103, USA; Department of Pathology & Translational Pathobiology, LSU Health, Shreveport LA 71103, USA; Department of Pathology and Laboratory Medicine Service, Overton Brooks VA Medical Center, Shreveport, Louisiana, USA

**Keywords:** Prostate cancer, NEPC, AdPCa, INSM1, Neuroendocrine differentiation, Lineage plasticity

## Abstract

Neuroendocrine prostate cancer (NEPC) is a highly aggressive and therapy-resistant subtype that arises from adenocarcinoma through lineage plasticity; however, the molecular mechanisms driving this transition remain incompletely defined. Insulinoma-associated protein 1 (INSM1), a zinc-finger transcription factor and established neuroendocrine lineage marker, has been implicated in a variety of neuroendocrine malignancies, yet its functional contribution to NEPC progression is not well understood. In this study, we demonstrate that INSM1 is consistently upregulated across NEPC patient tumors and experimental models, including both ASCL1⁺ and NEUROD1⁺ molecular subtypes, as revealed by integrated bulk and single-cell transcriptomic analyses. Functional studies revealed that INSM1 is sufficient to induce and necessary to maintain neuroendocrine lineage programs in prostate cancer, as overexpression promoted and depletion suppressed neuroendocrine-associated transcriptional networks. Mechanistically, pro-neural transcription factors, including ASCL1, NEUROD1, NEUROG3, and MYCN, directly or indirectly activate INSM1 expression, positioning it as a critical downstream effector of neuroendocrine lineage specification. Therapeutically, we identify homo-harringtonine (HHT), an FDA-approved protein synthesis inhibitor, as a potent suppressor of INSM1. HHT selectively reduces viability of INSM1-high NEPC cells at nanomolar concentrations, promotes ubiquitin-mediated degradation of INSM1, and significantly inhibits tumor growth *in vivo*. Notably, INSM1 depletion further enhances cellular sensitivity to HHT treatment. Collectively, our findings establish INSM1 as a key regulator of neuroendocrine plasticity and a promising therapeutic vulnerability in NEPC, providing a rationale for targeting INSM1 to suppress tumor progression.

## 1. Introduction

Neuroendocrine prostate cancer (NEPC) represents a rare but highly aggressive and clinically challenging subtype of prostate cancer that most commonly emerges because of therapeutic pressure. Prolonged androgen deprivation therapy (ADT) and treatment with second-generation androgen receptor (AR)-targeted agents (such as enzalutamide or abiraterone) can drive tumor evolution toward an AR-independent, neuroendocrine (NE) state (1, 2). This therapy-induced transformation reflects a fundamental shift in tumor cell identity rather than simple clonal selection. Clinically, NEPC is characterized by rapid disease progression, a propensity for visceral and lytic bone metastases, low or discordant prostate-specific antigen (PSA) levels, and minimal responsiveness to conventional AR-directed therapies. As a result, outcomes remain dismal, with reported median overall survival ranging from approximately 7 to 20 months despite aggressive multimodal treatment. These clinical features underscore an urgent unmet need for mechanistic insights and therapeutic strategies that target the biological processes driving this phenotypic transition. Among these, lineage plasticity-the ability of cancer cells to reprogram their differentiation state-has emerged as a central mechanism underlying resistance in advanced prostate cancer. Neuroendocrine differentiation (NED), a key manifestation of this plasticity, enables tumor cells to escape AR dependence by adopting transcriptional and epigenetic programs characteristic of neuronal and endocrine lineages (3, 4). Importantly, this transition is not merely a passive consequence of therapy but an active, regulated process involving coordinated changes in gene expression, chromatin accessibility, and signaling networks.

At the molecular level, NEPC is driven by a combination of recurrent genomic alterations and epigenetic reprogramming events. Loss of key tumor suppressors, including *RB1*, *TP53*, and *PTEN*, disrupts cell cycle control and genomic stability while facilitating de-differentiation (5). Concurrent amplification or activation of oncogenic drivers such as *MYCN* and *AURKA* further promotes the establishment of a neural-like transcriptional state (6). In parallel, epigenetic remodeling-mediated by chromatin modifiers and lineage-determining transcription factors-silences canonical AR signaling programs and activates neuronal and NE gene networks (7). Together, these alterations converge to rewire the transcriptional landscape, enabling prostate adenocarcinoma cells to trans-differentiate into an aggressive NE phenotype. Despite these advances, the key regulatory nodes that orchestrate this transcriptional shift remain incompletely defined. Identifying such nodes is essential for the development of therapeutic interventions aimed at preventing, delaying, or reversing NEPC evolution.

Insulinoma-associated protein 1 (INSM1) has emerged as a compelling candidate regulator in this context. INSM1 is a zinc-finger transcription factor that serves as a highly sensitive and specific nuclear marker of NE lineage across a wide spectrum of tumors. Functionally, INSM1 regulates the expression of genes involved in neuronal and endocrine differentiation, including *NEUROD1*, *YAP1*, and *HES1* (8–10). Mechanistically, INSM1 acts primarily as a transcriptional repressor through interactions with chromatin-modifying complexes, thereby shaping gene expression programs that define NE cell identity (10, 11). Its expression is tightly regulated during development and is largely restricted to embryonic and early postnatal stages in normal tissues, highlighting its role as a lineage-specific developmental factor. Beyond its diagnostic utility, INSM1 plays critical roles in developmental biology, where it governs key processes such as pancreatic endocrine differentiation, sympathoadrenal lineage specification, and neuronal precursor maturation (12–15). These functions position INSM1 at the intersection of multiple developmental pathways that are co-opted during tumorigenesis. Indeed, accumulating evidence from our group and others indicates that INSM1 is not merely a passive marker of NED but an active regulator of NED and tumor progression (8, 16–19). In several NE malignancies, including small cell lung cancer (SCLC) and neuroblastoma (NB), INSM1 has been shown to interact with core transcriptional networks and contribute to tumor cell survival and proliferation (20–22). To further interrogate its functional role, we previously developed an *INSM1* promoter-driven luciferase screening platform, which enabled the identification of small-molecule inhibitors capable of suppressing INSM1-associated signaling pathways (23). These studies provided functional evidence that INSM1 activity is druggable and contributes to the maintenance of NE tumor phenotypes across distinct cancer types. In the context of prostate cancer, our prior work demonstrates that INSM1 is robustly and specifically expressed in NEPC tissues and models (24, 25). These observations support a model in which INSM1 functions as a central regulatory hub linking upstream pro-neural transcription factors with downstream NE effector programs, thereby driving lineage plasticity and sustaining the NEPC phenotype. However, the precise mechanisms by which INSM1 coordinates these transcriptional networks, as well as its potential as a therapeutic target in prostate cancer, remain to be fully elucidated. In this study, we investigate the mechanistic role of INSM1 in NEPC progression, focusing on its position within the transcriptional circuitry governing NED and lineage plasticity. Furthermore, we evaluate the therapeutic potential of targeting INSM1-driven pathways as a strategy to disrupt NEPC evolution and improve outcomes in this lethal disease.

## 2. Results

### INSM1 is highly expressed in NEPC

We and others have previously demonstrated that INSM1 serves as a key NE tumor marker (16, 19). In prostate cancer, our data analysis reveals significant increased INSM1 *mRNA* levels in human NEPC relative to AdPCa in a compiled *RNA-seq* datasets derived from multiple studies (**Fig. 1A, B, red** arrow points to INSM1) (24, 26). Elevated INSM1 expressions are accompanied by increased levels of key NE-associated transcription factors, including ASCL1 and NEUROD1, as well as epigenetic regulators such as KDM1A (LSD1) and EZH2. Additionally, INSM1 expression positively correlates with established NE markers (SYP, CHGA, NCAM1) and NED-associated factors (ELAVL3, PCSK1, PCSK2), further supporting its role in the NEPC molecular program. Immuno-histochemistry (IHC) of INSM1, CHGA, and SYP revealed positive staining in human prostate cancer specimens. INSM1 expression was detected in 9/9 NEPC samples, compared with 1/40 adenocarcinoma samples. (**Fig. 1C**). These findings reinforce the role of INSM1 as a robust biomarker of NEPC and suggest its potential as a therapeutic target in this aggressive disease subtype. To investigate the cellular distribution of INSM1 in NEPC, we analyzed the Human Prostate Single-Cell Atlas (HuPSA), which integrates publicly available *single-cell RNA* sequencing data from 74 human prostate samples across six studies and contains 368,831 high-quality cells (25). Uniform Manifold Approximation and Projection (UMAP) analysis identified multiple cellular populations, including adenocarcinoma, NE, mesenchymal, and immune cell populations. Consistent with previous studies, NEPC cells could be further separated into two major subtypes characterized by expression of either ASCL1 or NEUROD1. These transcription factors displayed largely mutual exclusive expression patterns (25). In contrast, INSM1 expression was detected in both ASCL1⁺ and NEUROD1⁺ NE populations, indicating that INSM1 is not restricted to a specific NEPC subtype (**Fig. 1D**). INSM1 expression was also observed in normal NE cells, although these cells were present at low frequency compared with malignant NE populations. These findings demonstrate that INSM1 is broadly expressed across the major NE cell states identified in the prostate and is a shared feature of both ASCL1⁺ and NEUROD1⁺ NEPC subtypes.

**Figure 1.**
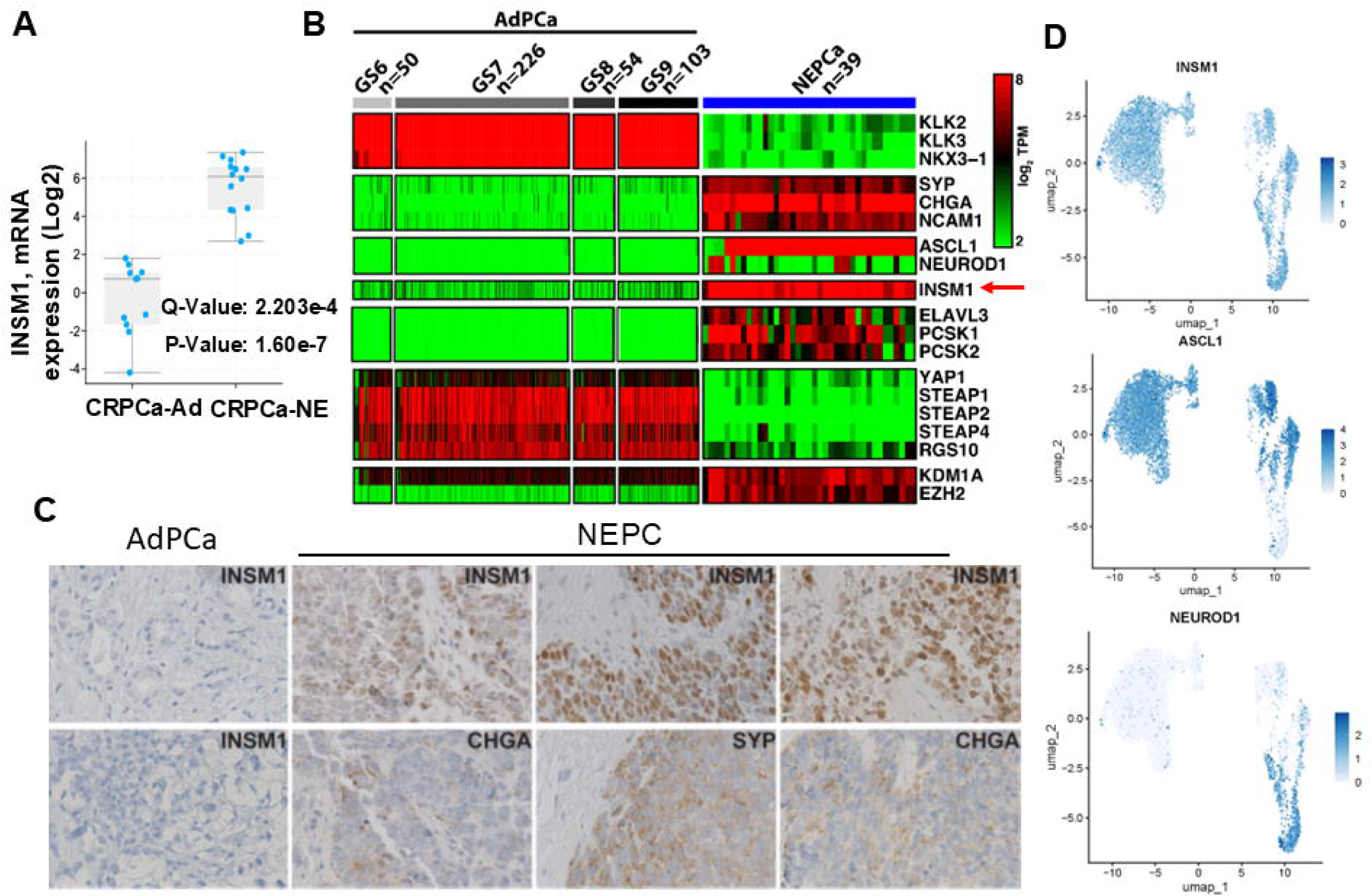
INSM1 is elevated in NEPC. **(A)** Data mining of INSM1 *mRNA* in Beltran NEPC dataset of cBioportal. **(B)** Data mining of compiled NEPC dataset in Human Prostate Single-Cell Atlas (HuPSA) demonstrates significantly increased INSM1 *mRNA* expression in human NEPC compared to AdPCa (pointed by red arrow). **(C)** IHC of INSM1, CHGA and SYP in human prostate cancer specimens. INSM1 expression was detected in 9/9 NEPC samples, compared with 1/40 adenocarcinoma samples. These data support INSM1 as a robust biomarker of NEPC. **(D)** INSM1 is expressed in both ASCL1⁺ and NEUROD1⁺ NEPC subtypes. *scRNA-seq* data was extracted from HuPSA dataset.

### INSM1 expression parallels NED marker program

To investigate lineage plasticity, we examined the capacity of prostate adenocarcinoma (AdPCa; luminal epithelial, AR-dependent) cells to transition toward alternative phenotypes, including basal-like, stem-like, and NE states. We utilized a panel of prostate cancer cell lines representing a continuum of differentiation and plasticity (**Fig. 2A**): LNCaP (AR⁺/NE⁻; early luminal), DU145 (AR⁻/NE⁻; lineage-switched non-NE), PC3 (AR⁻ with partial/weak NE features; progenitor/stem-like), NCI-H660 (AR⁻/strongly NE⁺; bona fide NEPC), and LASCPC-01 (AR⁻/low; mixed NE, a transitional plasticity model driven by lentiviral N-Myc and myristoylated-AKT1 expression) (27). To assess the relationship between INSM1 expression and NED, we analyzed *RNA-seq* data from the Broad Cancer Cell Line Encyclopedia (https://depmap.org/portal). Cell lines include non-cancerous prostate cells, prostate adenocarcinoma models, and prostate small cell carcinoma (H660) (**Fig. 2B**). INSM1 expression showed a strong positive correlation with canonical NE markers (CHGA, SYP, NCAM1) as well as NED-associated regulators (DLL3, MYCL, ASCL1). Consistent with this, INSM1 levels were markedly elevated in NEPC models (H660 and LASCPC-01), whereas AdPCa cell lines exhibited minimal expression. Notably, the INSM1 expression pattern closely mirrored that of established NE markers and NED-related transcriptional programs, supporting its association with NE lineage states.

**Figure 2.**
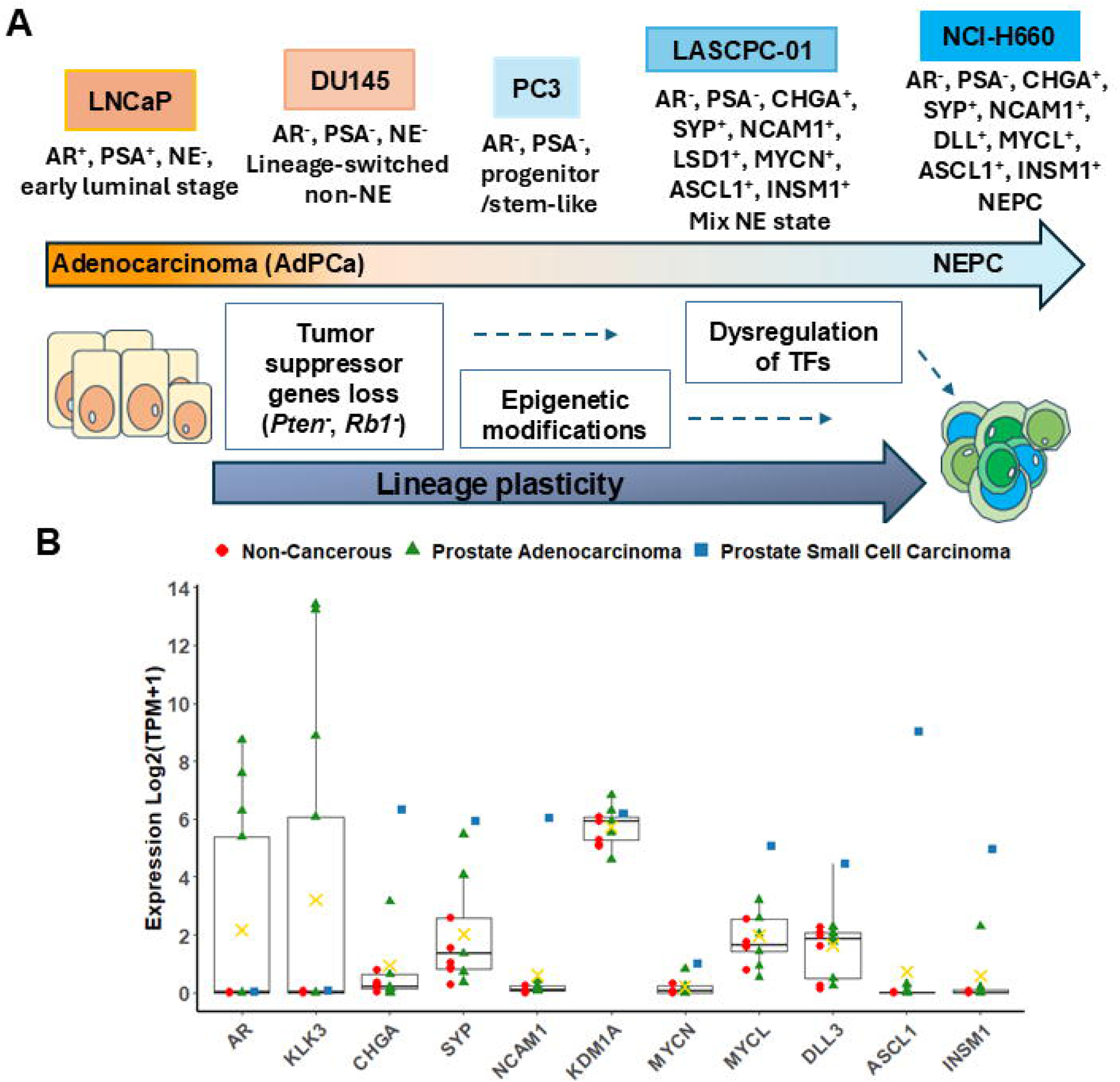
Lineage plasticity and the NE phenotype during prostate cancer trans-differentiation. **(A)** Multiple coordinated events contribute to the trans-differentiation of AdPCa to a NE phenotype, including loss of tumor suppressors (*PTEN* and *RB1*), epigenetic reprogramming, and dysregulation of key transcription factors. Prostate cancer cell lines, including LNCaP, DU145, PC3, LASCPC-01, and H660 represent distinct stages along this trans-differentiation continuum. INSM1 expression is predominantly elevated in NEPC cell lines (H660 and LASCPC-01), coinciding with increased expression of NE markers. **(B)** *RNA-seq* analysis derived from the Broad Cancer Cell Line Encyclopedia: https://depmap.org/portal revealed that H660 expresses high levels of INSM1, ASCL1, DLL3, MYCL, NCAM1, SYP, and CHGA (blue square).

### Transcriptomic analysis of INSM1 expression in prostate cancer

In LNCaP cells (AR-positive luminal adenocarcinoma), INSM1 overexpression (OE) robustly increased INSM1 transcript levels and triggered a coordinated transcriptional reprogramming (**Fig. 3A**). However, short-term INSM1 induction was insufficient to drive a complete NE phenotype in LNCaP cells. Heatmap shows log2 expression of genes involved in four different categories: NE lineage and neural-like (pink), AR/luminal signal (green), AR-driven mitotic regulator (yellow), and *p53*/stress/DNA damage response (orange). Gene were selected using differential expression thresholds of *log2FC>1, p < 0.001*. Several neural-associated genes, either upregulated (*SYT4, IFITM10*) or down regulated (*GAL, TMSB15A, BCYRN1*) in INSM1 overexpression, whereas canonical NE lineage regulators and markers were not detected in any changes. Instead, acute INSM1 overexpression (48-hour induction) predominantly suppressed the *AR/E2F1/FOXM1* proliferative axis, resulting in downregulation of key cell-cycle and mitotic regulators, including *E2F1, UBE2C, CDC20, CCNB1, FOXM1, MYBL2, E2F1, RRM2, CDK1, and TYMS*. Concurrently, INSM1 also inversely activated prostate carcinoma marker (*KLK3/PSA*), which against NEPC transition and markers associated with lipid metabolism (*AMACR, ACSL3, ELOVL5*). Genes related to the stress-response and growth-inhibitory pathways were evidenced by increased expression of *CDKN1A, MDM2, GADD45A, PMAIP1, PHLDA3, and HMOX1*. These transcriptional changes are consistent with cell-cycle arrest and the acquisition of partial lineage plasticity. Together, these findings suggest that INSM1 functions as an early facilitator of treatment-induced NE trans-differentiation by weakening the AR-dependent proliferative program and suppressing AR-driven mitotic regulators.

**Figure 3.**
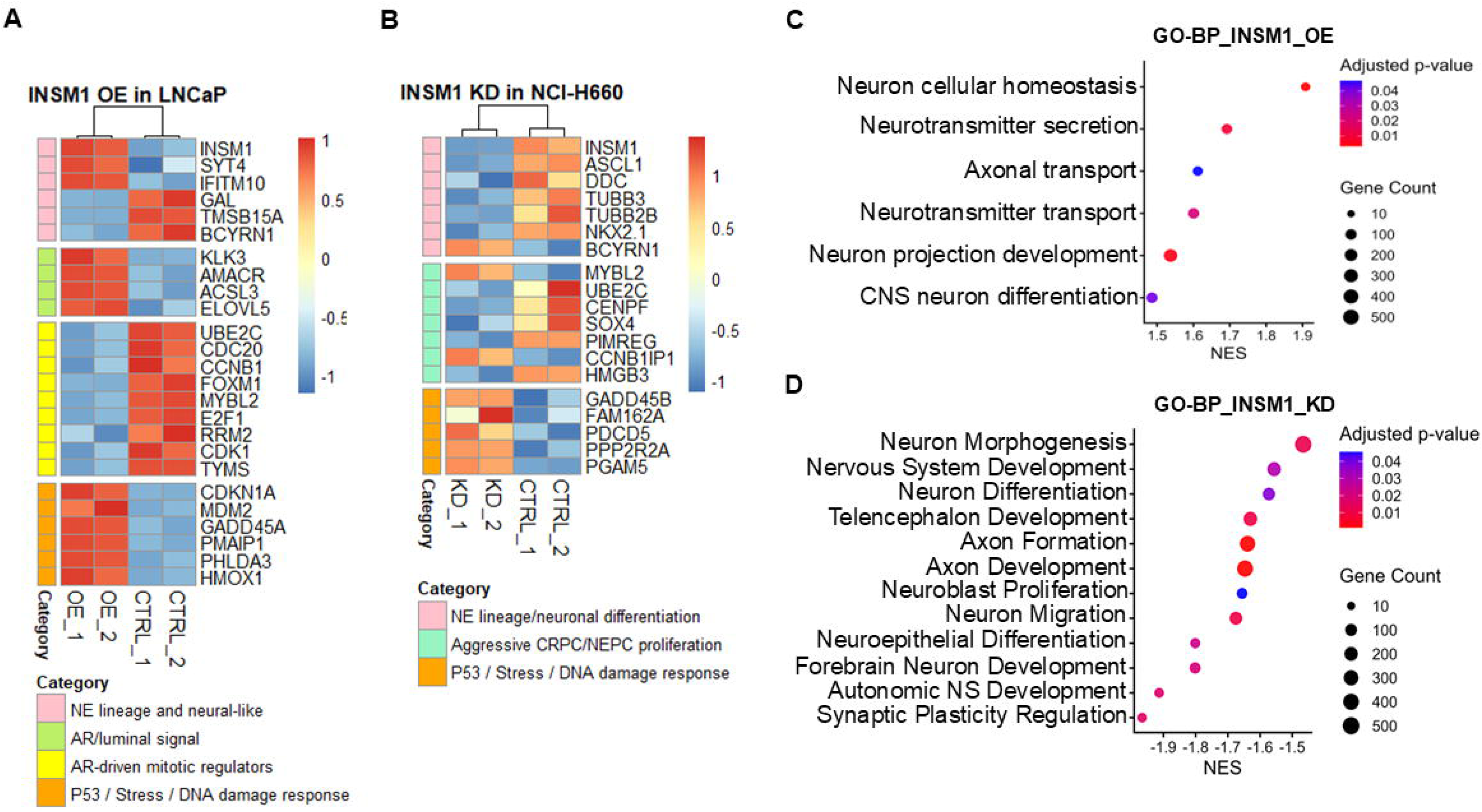
Transcriptomic analysis of INSM1 expression in prostate cancer. **(A)** Gene-expression profile from LNCaP cells following INSM1 overexpression. Heatmap shows log2 expression of genes involved in four different categories; NE lineage and neural-like (pink), AR/luminal signal (green), AR-driven mitotic regulator (yellow), and p53/stress/DNA damage response (orange). Color scale represents relative expression from low (blue) to high (red). Gene were selected using differential expression thresholds of *log2FC>1, p < 0.001*. **(B)** Gene-expression profile from NCI-H660 (NEPC) cells following INSM1 knockdown. Heatmap shows log2 expression of genes involved in three different categories; NE lineage/neuronal differentiation (pink), aggressive CRPC/NEPC proliferation (green), and p53/stress/DNA damage response (orange). Gene were selected using differential expression thresholds of *log2FC>1, p < 0.05*. Neuron-associated GO-BP enrichment analysis of *RNA-seq* from INSM1 overexpression **(C)** and knockdown **(D)** cells identified significant enrichment of neuronal development and differentiation pathways following INSM1 alteration. Pathways were filtered using *FDR < 0.05* and reduced by semantic similarity clustering (cutoff = 0.6).

In NEPC cells (NCI-H660), INSM1 functions as a master NE lineage factor that is required to maintain the NE transcriptional program. Heatmap shows log2 expression of genes involved in three different categories: NE lineage/neuronal differentiation (pink), aggressive CRPC/NEPC proliferation (green), and *p53*/stress/DNA damage response (orange). Gene were selected using differential expression thresholds of *log2FC>1, p < 0.05*. INSM1 knockdown reduced the expression of key NE regulators and markers, including *ASCL1, DDC, TUBB3, TUBB2B, and NKX2-1*. Furthermore, INSM1 was essential for sustaining the aggressive growth phenotype of NEPC cells. INSM1 depletion either suppressed (*UBE2C, CENPF, SOX4, PIMREG, HMGB3*) or activated (*BCYRN1, MYBL2, CCNB1IP1, GADD45B, FAM162A, PDCD5, PPP2R2A, and PGAM5*) in multiple *p53*/stress/DNA damage response and proliferation-associated genes implicated in CRPC/NEPC progression (**Fig. 3B**). Collectively, the *RNA-seq* analyses and orthogonal perturbation studies reveal distinct stage-specific functions of INSM1 during NEPC development. In adenocarcinoma cells, INSM1 acts as an early facilitator of the AdPCa-to-CRPC/NEPC transition by repressing the *AR/E2F1*-driven proliferative network while activating programs associated with NED, secretory architecture, and intracellular trafficking, thereby promoting cellular plasticity. This transcriptional pattern is accompanied by suppression of cell-cycle and translational machinery and closely mirrors the lineage switch observed in patient tumors and preclinical models. In contrast, once the NEPC state is established, tumor cells become dependent on INSM1 to maintain NE lineage identity and sustain aggressive tumor growth. Together, these findings identify INSM1 as both a key initiator of lineage reprogramming and a critical dependency factor required for the maintenance and progression of advanced CRPC/NEPC.

To further investigate the role of INSM1 in NE plasticity, we performed GSEA of GO Biological Process pathways following INSM1 overexpression in LNCaP cells and INSM1 knockdown in H660 NEPC cells. As shown in **Fig. 3C, D**, INSM1 overexpression enriched pathways associated with neuronal function, including neurotransmitter transport, neurotransmitter secretion, axonal transport, neuron projection regulation, and neuronal homeostasis, consistent with activation of NE lineage-associated programs. In H660 cells, INSM1 knockdown altered pathways involved in neuronal differentiation, axonogenesis, axon development, neuron migration, and nervous system development, indicating disruption of neuronal developmental programs. Together, these findings support a role for INSM1 in regulating transcriptional networks associated with NE identity and plasticity in prostate cancer.

### Functional role of INSM1 in NEPC

INSM1 was ectopically expressed in AdPCa cell lines representing AR⁺ (LNCaP) and AR⁻ (PC3) backgrounds. Overexpression of INSM1, like ASCL1, induced the expression of multiple NEPC-associated markers, including CHGA, SYP, and LSD1, while suppressing YAP1 expression (**Fig. 4C, D**). Conversely, knockdown of INSM1 using *Ad-INSM1-siRNA-6* versus *sc-siRNA* (scrambled) control in NEPC cells (LASCPC-01 and H660) resulted in decreased expression of key NEPC markers, including N-Myc, SYP, CHGA, NCAM1, and LSD1 (**Fig. 4A, B**). Overexpression or knockdown of INSM1 reflects either direct or indirect functional effect on target genes expression. Collectively, these results demonstrate that INSM1 is both sufficient to promote and necessary to maintain the NE phenotype in prostate cancer, supporting its critical role in lineage plasticity and NEPC trans-differentiation.

**Figure 4.**
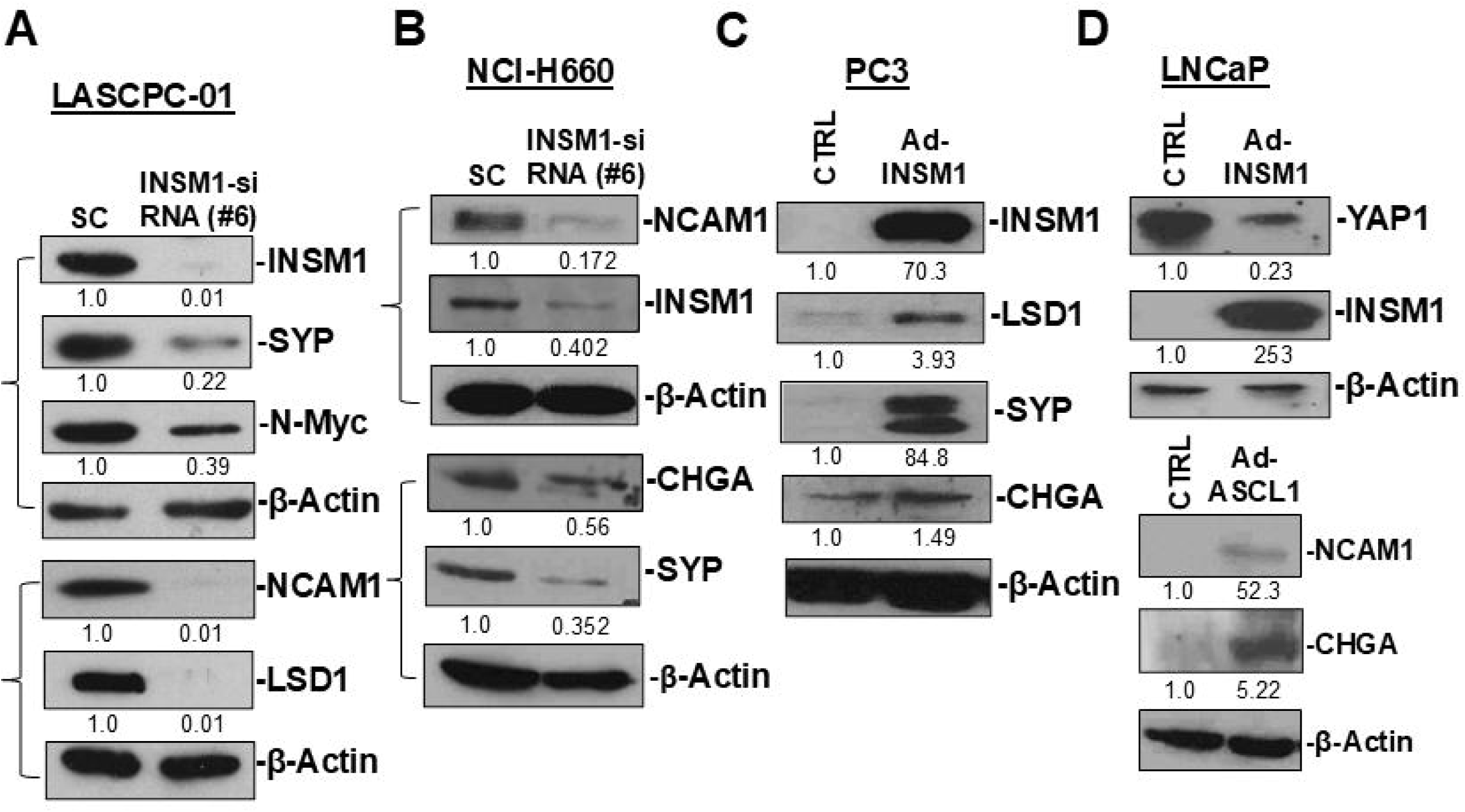
INSM1 modulates NE biomarker expression in prostate cancer cells. (A, **B)** Knockdown (KD) of INSM1 in NEPC cell lines (LASCPC-01 and H660) reduces the expression of key NE markers and associated factors, including N-Myc, SYP, NCAM1, CHGA, and LSD1. **(C)** Overexpression of INSM1 in PC3 cells induces the expression of NE markers, including LSD1, SYP, and CHGA. **(D)** Overexpression of INSM1 in LNCaP cells represses YAP1 expression while overexpression of ASCL1 inducing NCAM1 and CHGA. Protein levels were normalized to β-actin and are presented as relative expression ratios.

### Regulation of INSM1 expressions

The tightly controlled activity of the *INSM1*-promoter restricts its expression largely to NE tumors, including NEPC. This restricted expression pattern prompted us to investigate the molecular mechanisms governing *INSM1*-promoter activation. We investigate the transcriptional regulatory network in prostate cancer. In AdPCa cell lines (LNCaP, DU145, and PC3), adenoviral-mediated expressions of NEUROG3, NEUROD1, ASCL1, or N-Myc reproducibly induced INSM1 expression (**Fig. 5**). These data support a conserved transcriptional regulatory mechanism controlling INSM1 expression across NE cell types and tumor lineages, highlighting its central role in NED programs.

**Figure 5.**
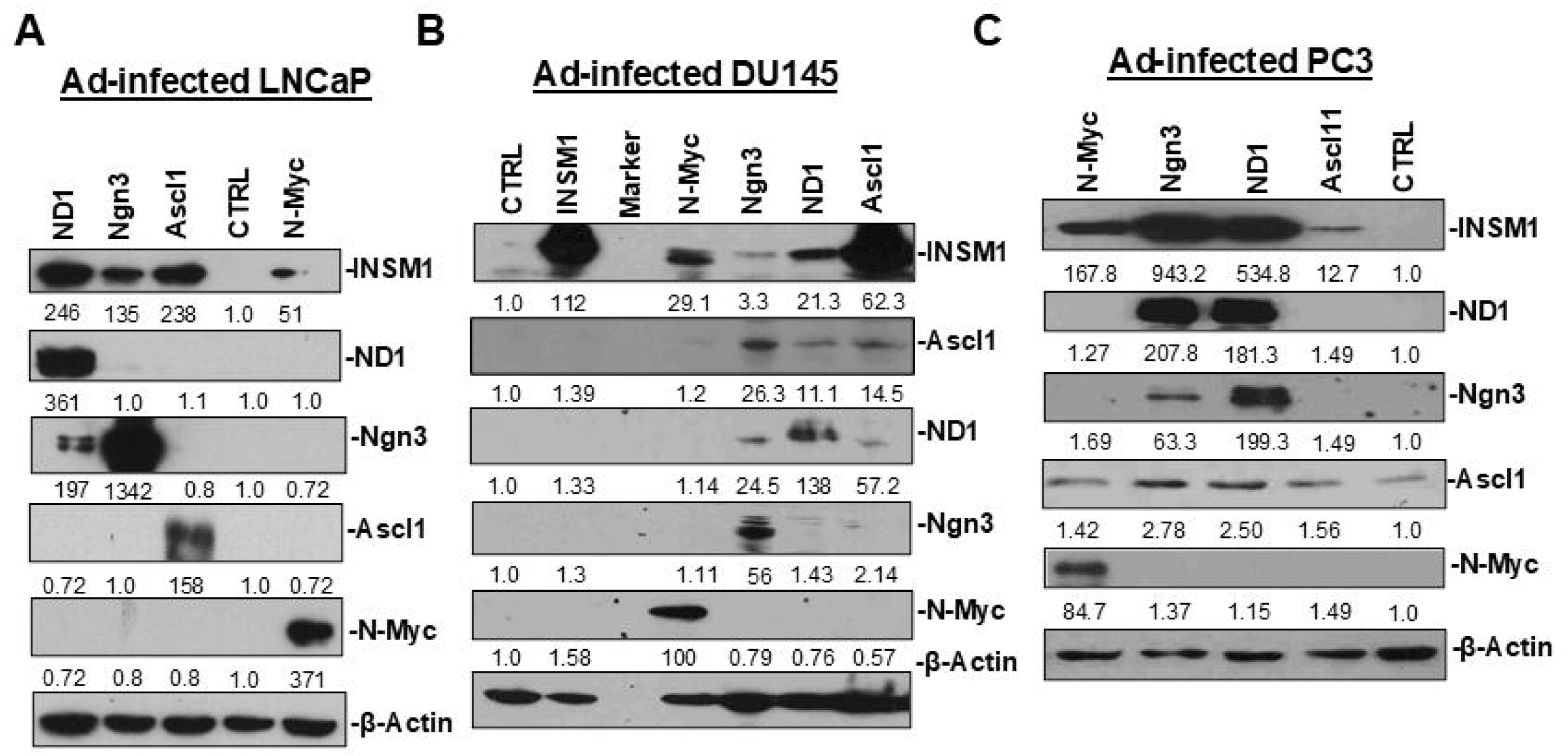
Pro-neural transcription factors induce INSM1 expression in prostate cancer cells. Pro-neural factors ASCL1, NEUROD1, NEUROG3 (Ngn3), and N-Myc each induce INSM1 expression. **(A)** LNCaP, **(B)** DU145, and **(C)** PC3 cells were infected with adenoviral vectors encoding these factors (Ad-ND1, Ad-Ngn3, Ad-Ascl1, or Ad-N-Myc) for 5 days. Expression of exogenous transcription factors and endogenous INSM1 was assessed by Western blot analysis. Protein levels were normalized to β-actin and are presented as relative expression ratios.

### INSM1 as a therapeutic target

To evaluate the therapeutic potential of INSM1 in NEPC, we examined the effects of INSM1 knockdown on tumor cell survival and growth. Silencing INSM1 using *Ad-INSM1-siRNA-6* in LASCPC-01 cells significantly reduced cell viability and tumor growth compared to control *Ad-sc-siRNA*-treated cells (**Fig. 6A, B**), indicating that INSM1 expression is critical for NEPC cell survival and tumor progression. These findings suggest that targeting INSM1 may represent an effective therapeutic strategy for NEPC. A recent study using DKO (*Pten^null^*/*Rb1^null^*) murine models identified early transitioning NE cell populations consisting of ASCL1⁺ and POU2F3⁺ clusters, both of which expressed INSM1 (5). Together, these data demonstrate that INSM1 is expressed not only in fully developed NEPC cells but also in intermediate cell populations undergoing NED. Consistent with this model, overexpression of INSM1 induces NEPC-associated biomarkers, whereas INSM1 knockdown suppresses their expressions. Notch signaling HES1 negatively regulates INSM1, while YAP1 negatively regulates Wnt/b-catenin signaling, which is critically important for N-Myc activation of INSM1 (1, 8, 21). Significantly, multiple oncogenic driver pathways implicated in NEPC converge to activate INSM1 expression (**Fig. 6C**). This central positioning working model supports INSM1 as a unique and attractive therapeutic target, whereas its inhibition may simultaneously disrupt multiple NEPC-driving pathways. Together, these results support INSM1 as a promising target for therapeutic intervention in NEPC.

**Figure 6.**
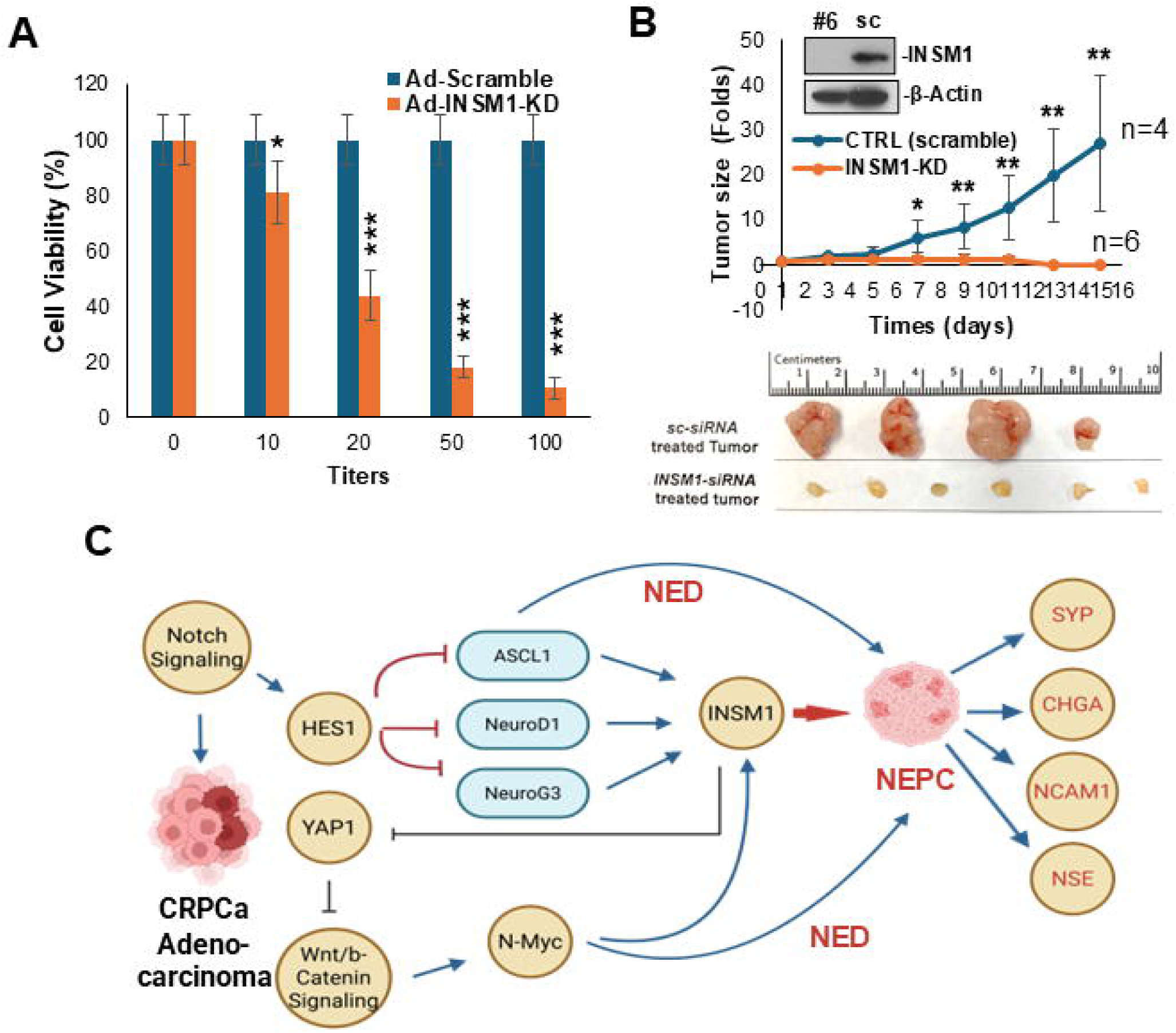
INSM1 knockdown suppresses NEPC cell viability and tumor growth that illustrates a working model of NEPC trans-differentiation. (A) Cell viability of LASCPC-01 cells following treatment with increasing titers of *INSM1-siRNA (#6)* or control *scrambled siRNA* (*sc-siRNA*) for 2 days. (B) LASCPC-01 cells (5 × 10⁶) infected with *INSM1-siRNA-6* or *sc-siRNA* were injected subcutaneously into the flanks of nude mice, and tumor growth was monitored. Western blot analysis confirms efficient knockdown (KD) of INSM1. (C) Proposed *<u>working model</u>* of AdPCa trans-differentiation into NEPC. ADT in castration-resistant prostate cancer (CRPCa) promotes lineage plasticity and the development of ADT treatment-related NEPC (tNEPC). Pro-neural transcription factors (ASCL1, NEUROD1, NEUROG3) and N-Myc induce INSM1 and NE marker expressions. These pathways are proposed to converge on INSM1, which in turn represses YAP1 expression. Reduced YAP1 activity leads to activation of Wnt/β-catenin signaling, promoting MYC family gene expressions and establishing a feed-forward loop that sustains INSM1 activation.

### Pro-neural transcription factors induce INSM1

To identify potential direct targets of INSM1 during NED, we integrated known genes with publicly available INSM1 *ChIP-seq* data derived from murine pancreatic β cells, due to the lack of human INSM1 *ChIP-seq* dataset. *ATAC-seq* profiles in LNCaP and H660 cells highlight differences in chromatin accessibility (**Fig. 7A**). *ChIP-seq* analysis of ASCL1 and NEUROD1 in SCLC cells. Alignment of the human genome to *ATAC-seq* peaks in LNCaP and H660 cells reveals that ASCL1 and NEUROD1 binding sites are enriched in chromatin regions that are more accessible in H660 cells (**Fig. 7B, C**). The integrative analysis identified candidate INSM1 target genes that are either upregulated or downregulated in NEPC relative to AdPCa (**Fig. 7D–H**). Notably, upregulated targets include NE-associated genes such as *ELAVL3*, *ELAVL4* (28), and *PCSK1/PCSK2* (29), whereas downregulated targets include *YAP1* (*9*), *STEAP1/2/4* (*30*), and *RGS10* (*31*). Pro-neural transcription factors, including ASCL1, NEUROD1, NEUROG3, N-Myc, and L-Myc (32), are highly expressed in human NEPC specimens and NEPC cell models such as H660. ASCL1- and NEUROD1-driven programs define the major molecular subtypes of NEPC (33–35), with INSM1 expressed across both subtypes (**Fig. 1**). Consistent with this, our experimental data demonstrates that overexpression of ASCL1, NEUROD1, NEUROG3, or N-Myc robustly induces INSM1 expression in prostate cancer cells (**Fig. 5**). Furthermore, recent studies have established INSM1 as a direct transcriptional target of ASCL1 (22). Supporting this, our analysis of publicly available *ChIP-seq* datasets reveals direct binding of ASCL1 and NEUROD1 at the INSM1 promoter region (**Fig. 7C**). These findings demonstrate that pro-neural transcription factors directly regulate INSM1 expression, placing INSM1 as a central downstream effector within the NE transcriptional network in NEPC.

**Figure 7.**
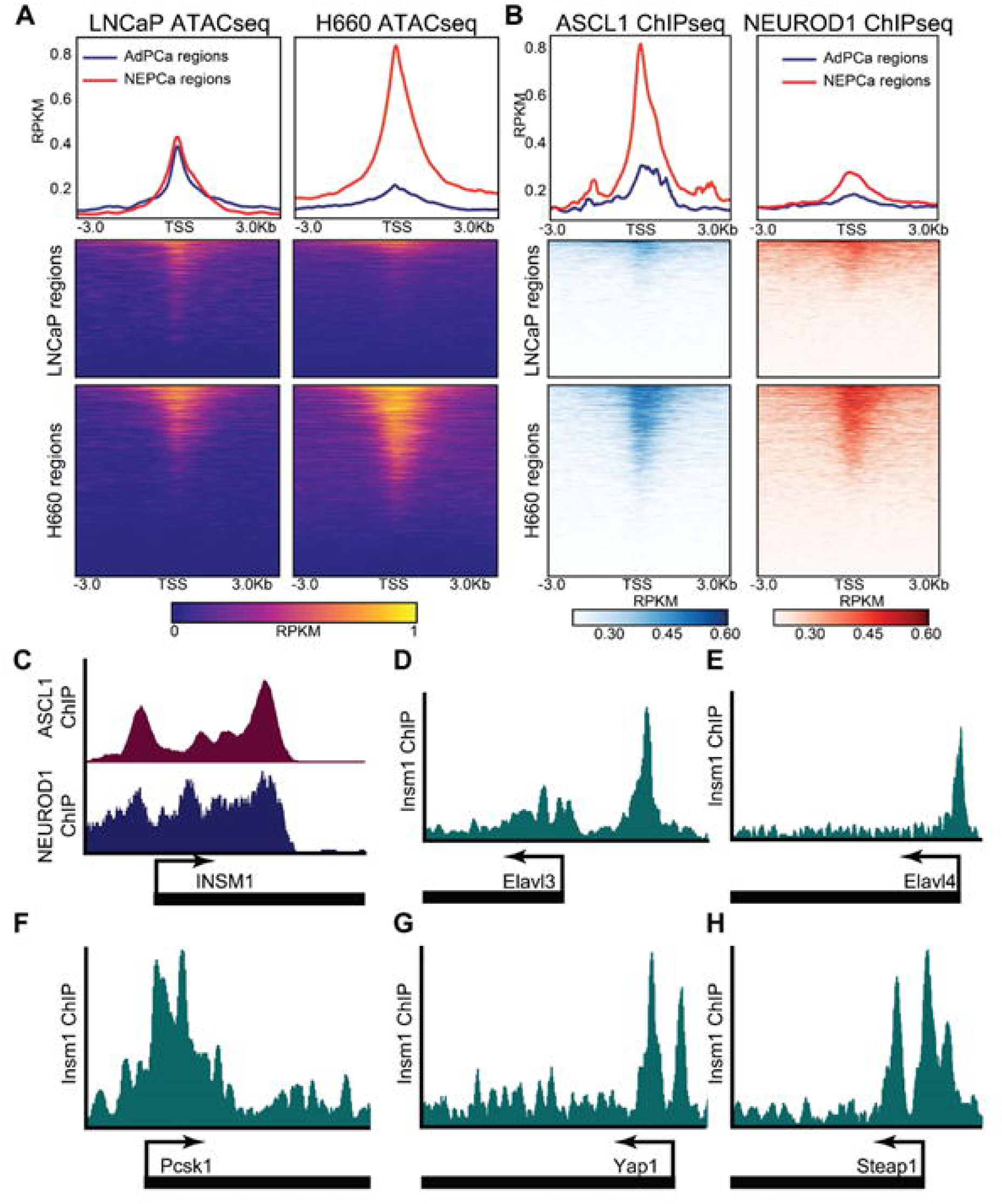
Integrated analysis of publicly available chromatin accessibility and *ChIP-seq* datasets. **(A)** *ATAC-seq* profiles in LNCaP and H660 cells highlight differences in chromatin accessibility. **(B)** *ChIP-seq* analysis of ASCL1 and NEUROD1 in SCLC cells. Alignment of the human genome to *ATAC-seq* peaks in LNCaP and H660 cells reveals that ASCL1 and NEUROD1 binding sites are enriched in chromatin regions that are more accessible in H660 cells. **(C)** *ChIP-seq* tracks showing ASCL1 and NEUROD1 binding at the *INSM1*-promoter region. **(D–H)** Analysis of publicly available *ChIP-seq* datasets identifies INSM1 binding at the promoters of candidate target genes, including *ELAVL3/4*, *PCSK1*, *YAP1*, and *STEAP1*.

### Anti-cancer efficacy of INSM1 inhibitor in NEPC

Accumulating evidence supports a central role for INSM1 in NED, highlighting it as a promising therapeutic target in NEPC. In our previous studies, screening of small-molecule libraries in NB cells identified several compounds capable of inhibiting INSM1 expression such as homo-harringtonine (HHT; *IC₅₀* = 0.034 μM) (22, 36). In the current study, we evaluated the therapeutic efficacy of HHT in NEPC and prostate cancer models. Comparative *IC₅₀* analysis across prostate cancer cell lines (DU145, LN95, PC3, LNCaP, H660, and LASCPC-01) revealed that HHT exhibits markedly greater potency in NEPC cells (nanomolar range) than in AdPCa cells (micromolar range) (**Fig. 8A**). Notably, LASCPC-01 cells displayed approximately 10-fold greater sensitivity to HHT than H660 cells (*IC₅₀*: 3 nM vs. 30 nM), consistent with their higher levels of INSM1 expression by Western blot analysis (**Fig. 8B**). HHT treatment also produced a dose-dependent reduction in LASCPC-01 cell viability (**Fig. 8C**). At the molecular level, HHT treatment reduced the expression of INSM1 and key NEPC biomarkers, including LSD1, NCAM1, SYP, and NSE (**Fig. 8D**). Concurrently, HHT increased levels of H3K4me2, a known target of LSD1 and upregulated *TP53* expression, suggesting that HHT may exert its effects through modulation of histone methylation and activation of tumor suppressor pathways. Importantly, HHT treatment significantly inhibited tumor growth *in vivo* (**Fig. 8E**). Furthermore, INSM1 knockdown enhanced the sensitivity of NEPC cells (LASCPC-01, H660) to HHT, resulting in a 3–6-fold reduction in *IC₅₀* values (**Fig. 9**), indicating an additive interaction between INSM1 suppression and HHT treatment. HHT (omacetaxine mepesuccinate), a semi-synthetic derivative with improved bioavailability, has been approved by the U.S. FDA for the treatment of chronic myeloid leukemia (CML) (37), with established pharmacokinetics and safety profiles in patients with advanced acute myeloid leukemia (AML) (38, 39). These clinical data support the feasibility of repurposing HHT as a therapeutic agent for NEPC. Notably, HHT demonstrates enhanced efficacy in INSM1-expressing NEPC cells, suggesting selective targeting of NE tumor populations. Targeting INSM1 during the transition from AdPCa to NEPC, particularly in combination with ADT may disrupt lineage plasticity and create a less permissive tumor microenvironment for NED. These findings highlight the therapeutic potential of targeting INSM1, with HHT representing a promising candidate for the treatment of NEPC with strong translational relevance.

**Figure 8.**
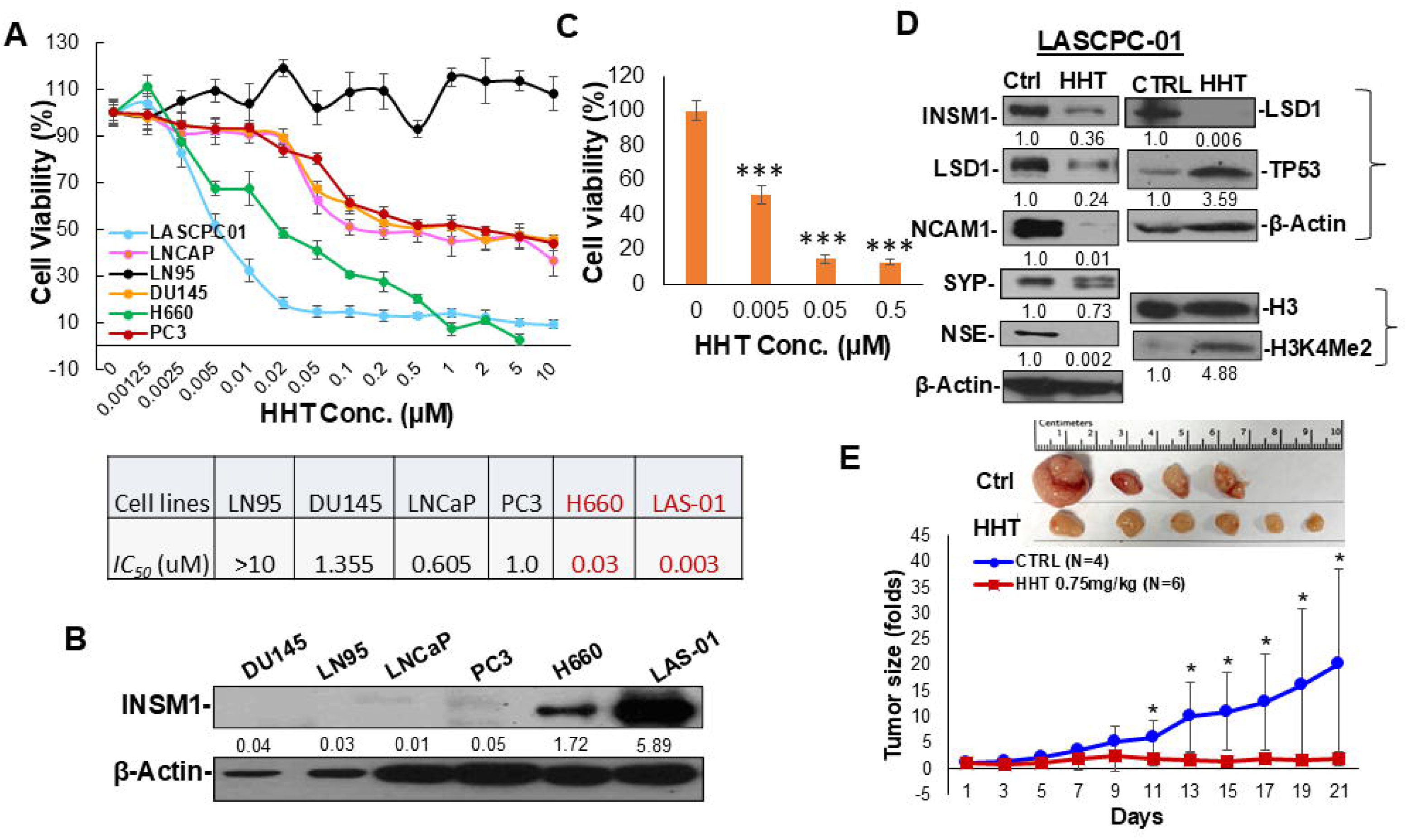
Effects of HHT on NEPC cells. **(A)** *IC₅₀* values of the INSM1 inhibitor HHT across prostate cancer cell lines, including non-NEPC and NEPC models. Cell viability assays were used to determine *IC₅₀* values, as summarized in the table. **(B)** Western blot analysis showing INSM1 protein expression across the indicated cell lines. Protein levels were normalized to β-actin and are presented as relative expression ratios. **(C)** Dose-dependent effects of HHT on LASCPC-01 cell viability. **(D)** Western blot analysis demonstrates that HHT treatment reduces the expression of INSM1 and NEPC-associated markers (LSD1, NCAM1, SYP, and NSE), while increasing H3K4me2 and TP53 levels in LASCPC-01 cells. **(E)** *In vivo* tumor growth assay in which LASCPC-01 cells (5 × 10⁶) were injected subcutaneously into nude mice, followed by daily treatment with HHT (0.75 mg/kg) for 21 days. Tumor growth was significantly inhibited in the HHT-treated group (*t test, *p < 0.05*).

**Figure 9.**
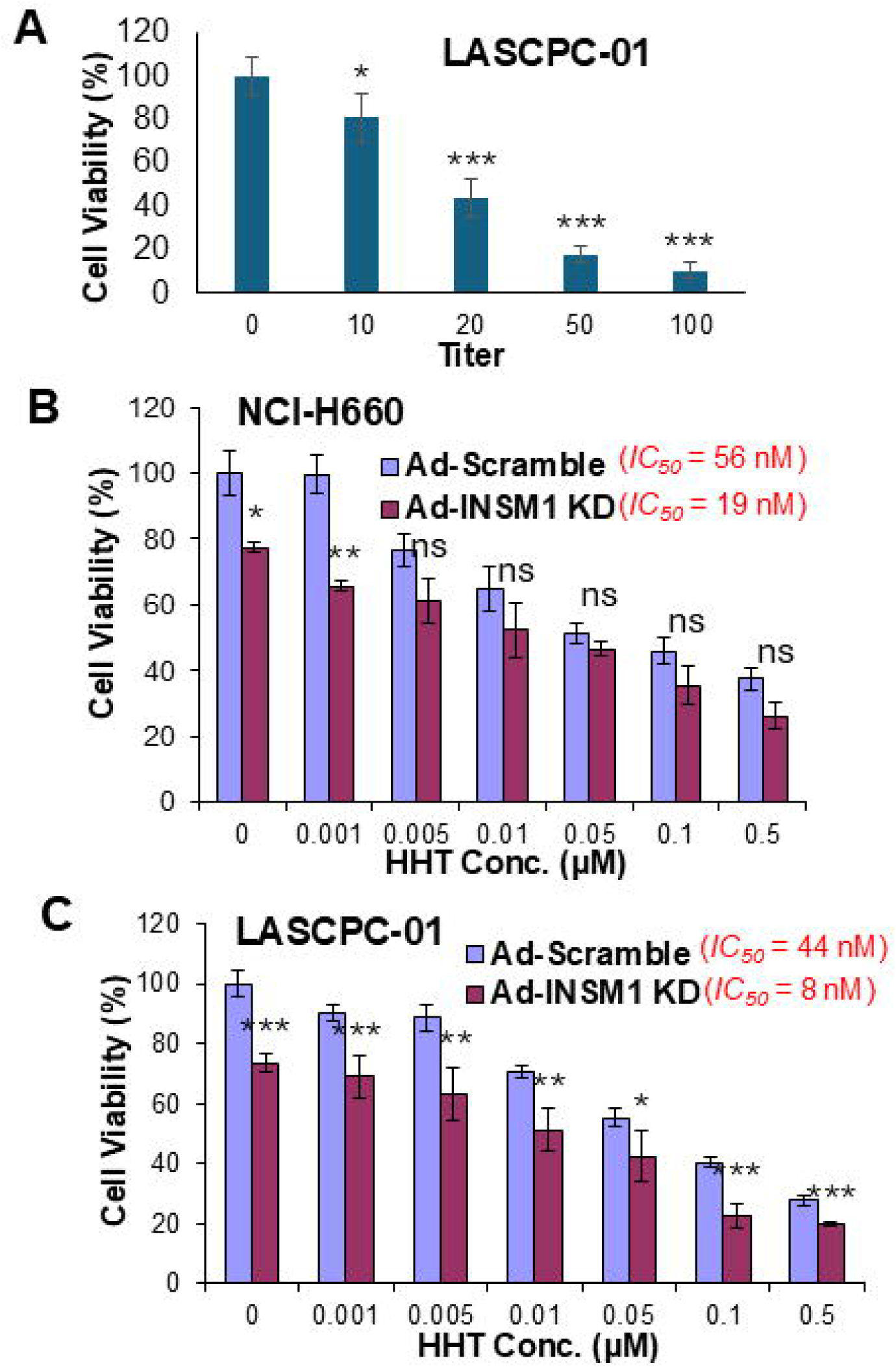
Combined effects of INSM1 knockdown (KD) and HHT on NEPC cells. **(A)** INSM1 knockdown using increasing titers of *Ad-INSM1-siRNA* results in progressive suppression of NEPC cell viability. INSM1 depletion enhances the cytotoxic effects of HHT. Comparison of single-agent treatment (HHT alone) versus combination treatment (INSM1 KD + HHT) demonstrates increased sensitivity (reduced *IC₅₀*) in NEPC cell lines, including **(B)** NCI-H660, **(C)** LASCPC-01. Statistical significance is indicated as follows: *\*p < 0.05, **p < 0.01, ***p < 0.001*; ns, not significant.

### HHT destabilizes INSM1 protein in NEPC cells

HHT is known to inhibit the initial elongation step of protein synthesis, suggesting that it may reduce the stability and half-life (t₁/₂) of INSM1 protein. Consistent with this, HHT treatment in NEPC cells (LASCPC-01) markedly decreased INSM1 protein levels. However, in the presence of the proteasome inhibitor MG132, INSM1 levels remained relatively stable regardless of HHT treatment (**Fig. 10A**), indicating that INSM1 degradation occurs through a proteasome-dependent mechanism. We next assessed INSM1 protein stability using a cycloheximide (CHX) chase assay (20 µg/mL) over a 0–12 h time course (**Fig. 10B**). Under basal conditions, INSM1 exhibited a half-life of approximately 3.997 hours. Treatment with HHT (100 nM) reduced the half-life to ∼2.502 hours, while combined treatment with CHX and HHT further accelerated INSM1 degradation, reducing the half-life to ∼1.554 hours (**Fig. 10C**). These findings indicate that ongoing protein synthesis is required to maintain INSM1 levels and that HHT accelerates INSM1 turnover via the ubiquitin–proteasome pathway. Taken together, these results demonstrate that regulation of INSM1 protein stability plays a critical role in maintaining NEPC tumor progression.

**Figure 10.**
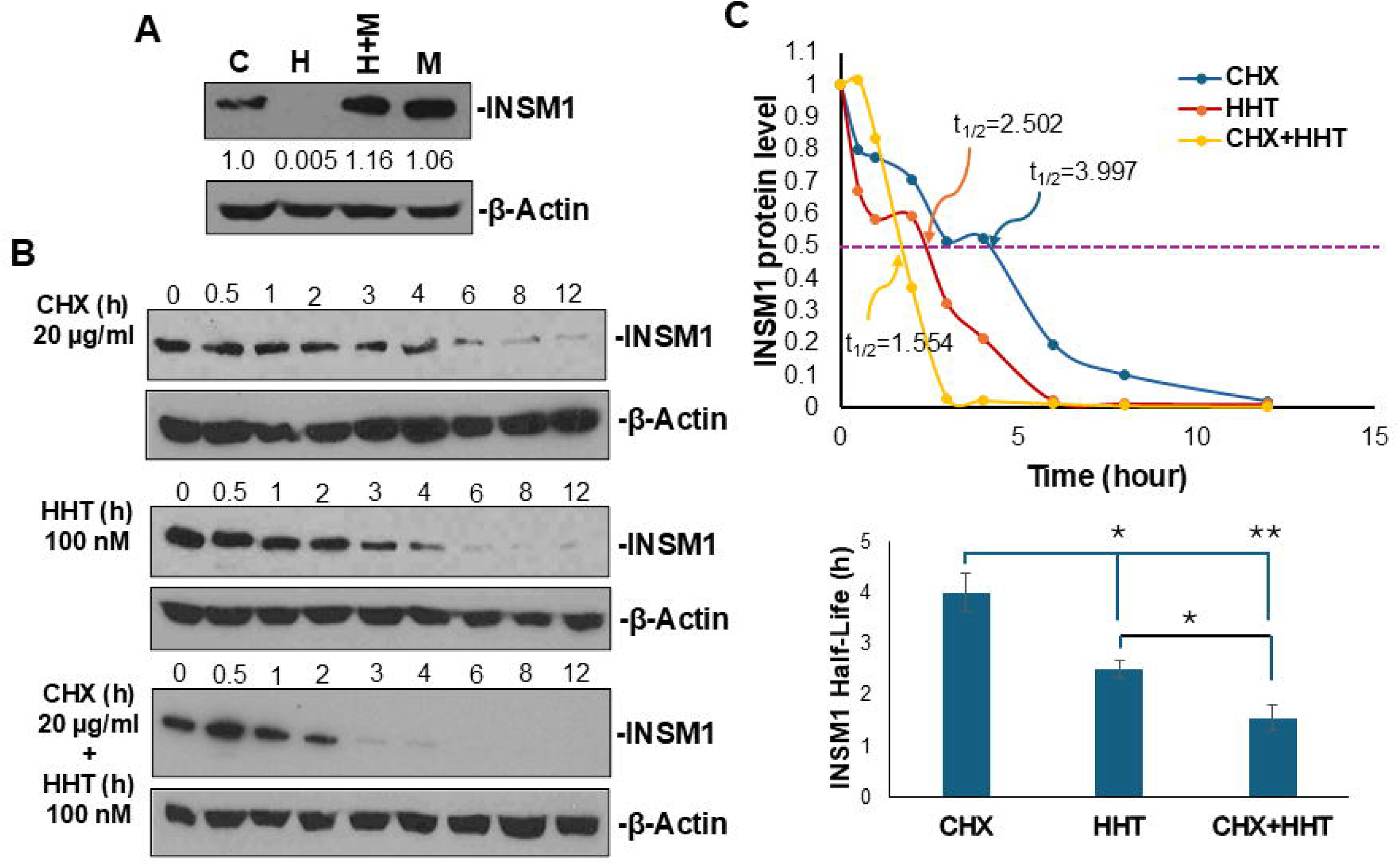
HHT promotes ubiquitin-mediated degradation and reduces stability of INSM1 protein. **(A)** Western blot analysis showing that HHT treatment markedly reduces INSM1 protein levels in LASCPC-01 cells, whereas co-treatment with the proteasome inhibitor MG132 maintains INSM1 levels, indicating proteasome-dependent regulation. **(B)** Cycloheximide (CHX; 20 µg/mL) chase assay (0–12 h) assessing INSM1 protein stability under control and HHT-treated conditions. **(C)** Quantification of INSM1 protein half-life (*t₁/₂*) showing accelerated degradation with HHT treatment (*t₁/₂* ≈ 2.502 h) compared to control (t₁/₂ ≈ 3.997 h), with further reduction upon combined CHX and HHT treatment (*t₁/₂* ≈ 1.554 h). These results indicate that HHT enhances INSM1 turnover through the ubiquitin–proteasome pathway.

## 3. Discussion

Neuroendocrine prostate cancer (NEPC) remains a highly aggressive and treatment-refractory malignancy, characterized by poor clinical outcomes and a lack of effective targeted therapies. A defining feature of NEPC is its capacity for lineage plasticity under therapeutic pressure, whereby AdPCa cells lose AR-dependence and adopt a NE phenotype that is largely unresponsive to standard-of-care treatments. This phenotypic transition highlights a critical unmet need to identify central regulators of NE lineage commitment that can function both as actionable therapeutic targets and clinically informative biomarkers. In this context, INSM1 emerges as a compelling candidate due to its strong specificity for the NE lineage and its functional involvement in NEPC biology.

During NEPC progression, neural and endocrine gene expression programs are activated while prostate lineage-specific differentiation pathways are suppressed (40). Many upstream NE transcriptional regulators, including ASCL1, NEUROD1, NEUROG3, and MYC family members are characterized by rapid protein turnover, suggesting that sustained protein synthesis may be essential to maintain NE identity. Consistent with this, our findings demonstrate that INSM1 is a highly dynamic protein whose levels decline rapidly upon translational inhibition with homo-harringtonine (HHT; omacetaxine mepesuccinate). This intrinsic instability indicates a dependence on continuous translation, revealing a potential therapeutic vulnerability that can be exploited through pharmacologic targeting of protein synthesis.

Clinically, NEPC is associated with a median survival of less than one year, marked by diminished AR signaling, resistance to ADT, and elevated expression of canonical NE markers, including CHGA, SYP, and INSM1 (41–43). While *de novo* NEPC is rare (<2% of prostate cancers), therapy-induced NEPC (t-NEPC) arises in more than 20% of patients with metastatic CRPC, driven by prolonged exposure to ADT and next-generation AR pathway inhibitors (1, 44, 45). Despite increasing knowledge of the genomic and transcriptional landscape of NEPC, the molecular mechanisms governing the transition from CRPC to NEPC remain incompletely defined, limiting the development of effective targeted interventions. Our data position INSM1 as a central regulatory node that links upstream pro-neural transcription factors (e.g., ASCL1, NEUROD1, NEUROG3, and MYCN) to downstream NE effector genes such as CHGA, SYP, and NCAM1 (10, 12). This role is consistent with prior observations in developmental systems, where these transcription factors drive INSM1 expression during endocrine and neuronal differentiation (13, 15). Accordingly, INSM1 functions as a conserved regulator of endocrine lineage specification and trans-differentiation across tissues (14, 46–48). In the context of cancer, our findings suggest that INSM1 contributes to the maintenance of progenitor-like cellular states and enhances phenotypic plasticity, thereby facilitating lineage reprogramming and tumor adaptation under therapeutic stress (49). Importantly, our results support a model in which INSM1 operates within a complex, non-linear regulatory network involving both feedforward and feedback interactions with other lineage-specifying factors. For instance, ASCL1-INSM1, NEUROG3-NEUROD1-INSM1, and N-Myc-INSM1 regulatory axes may reinforce NE lineage identity, while reciprocal interactions with factors such as YAP1 or HES1 may contribute to hybrid or intermediate cellular states. Systematic dissection of these regulatory circuits through targeted perturbation, individually and in combination, will be critical for understanding the mechanisms of lineage switching and identifying points of therapeutic intervention.

From a translational standpoint, NEPC remains a major clinical challenge, particularly with respect to early detection and prevention during ADT. Our preclinical models, which enable controlled induction of NED, provide a valuable platform for evaluating early intervention strategies. Integration of biomarker-driven monitoring approaches including INSM1 expression and circulating NE markers such as CHGA, Pro-GRP, and neuron-specific enolase (NSE) may improve patient stratification and optimize the timing of therapeutic interventions (50, 51). In parallel, emerging evidence suggests that INSM1 is linked to metabolic and epigenetic regulation. Alterations in methionine cycle metabolism, particularly in the balance of S-adenosylmethionine (SAM) and S-adenosylhomocysteine (SAH), have been shown to influence *INSM1* promoter activity and global histone methylation, implicating epigenetic remodeling as a key determinant of NE lineage commitment (49). These findings position INSM1 as a central integrator of transcriptional, metabolic, and epigenetic programs in NEPC.

Notably, our study demonstrates that HHT exerts potent anti-tumor activity in NEPC models, at least in part by targeting INSM1-dependent pathways. Although HHT is a well-established inhibitor of translational elongation and is clinically approved for certain hematologic malignancies (52, 53), our findings reveal a distinct functional impact in solid tumors. Specifically, HHT suppresses NED and tumor growth, with enhanced efficacy in INSM1-high contexts. This suggests that NEPC tumors with elevated INSM1 expression may be particularly reliant on continuous protein synthesis to sustain their lineage identity, thereby conferring selective vulnerability to translational inhibition.

In summary, this study establishes INSM1 as a central regulator of NE lineage plasticity and a key driver of NEPC progression. By elucidating its role within interconnected transcriptional, metabolic, and epigenetic networks, we provide a mechanistic framework for the development of novel therapeutic strategies. Targeting INSM1 directly, or through disruption of its regulatory dependencies, represents a promising avenue for intervention. These findings have important translational implications and may ultimately contribute to improved outcomes for patients with this highly lethal disease.

## 4. Materials and methods

### Cell culture, antibodies, and chemical reagents

Human prostate cancer cell lines LASCPC-01 (CVCL_UE17), NCI-H660 (CVCL_1576), LNCaP (CVCL_1379), LN95 (CVCL_ZC87), DU145 (CVCL_0105), and PC-3 (CVCL_0035) were obtained from ATCC (Manassas, VA, USA). All lines were authenticated by STR profiling, verified morphologically, used at <35 passages, and routinely tested for mycoplasma contamination. Cells were cultured in RPMI 1640, EMEM, or F-12K media supplemented with 10% fetal bovine serum and 1% penicillin/streptomycin, according to supplier guidelines, and maintained at 37 °C with 5% CO₂. Homo-harringtonine (HHT; Sigma-Aldrich), MG132 (Selleck), and cycloheximide (CHX; US Biological) were dissolved in DMSO; final DMSO concentrations did not exceed 0.1% (v/v). Primary antibodies included INSM1 (A8), N-Myc (B8.4B), β-actin, and p53 (Santa Cruz Biotechnology); LSD1, histone H3, and H3K4me2 (Cell Signaling Technology); CHGA (Protein-tech); ASCL1 (BD Biosciences); NEUROD1 and synaptophysin (Thermo-Fisher Scientific); and NEUROG3 and NCAM1 (DSHB). HRP-conjugated anti-mouse and anti-rabbit secondary antibodies were from Bio-Rad. Unless otherwise stated, antibodies were used at 1:1000 dilution.

### Immunohistochemistry (IHC) staining

A total of 49 human prostatic specimens were utilized in this study, obtained from Overton Brook VA medical Center, Louisiana State University Health Sciences Center at Shreveport Biorepository Core, and Tissue for Research, as described previously (54). Tissue sections (5-μm thick) were prepared from formalin fixed paraffin-embedded specimens, and IHC was performed following established protocols. The specimens were classified based on histology, Gleason grades, and the expression of NEPC biomarkers, CHGA, SYP, INSM1. All samples were collected and utilized in accordance with protocols approved by the Institutional Review Board of LSU Health Shreveport.

### Western blot analysis

Cells were lysed in buffer containing 10 mM Tris–HCl (pH 7.5), 150 mM NaCl, 10% glycerol, 1% Triton X-100, 1 mM DTT, 0.2 mM PMSF, protease inhibitors (aprotinin, leupeptin), and phosphatase inhibitors (Na₃VO₄, NaF). Equal protein amounts (100 μg) were resolved by SDS–PAGE and transferred to nitrocellulose membranes (Bio-Rad). Membranes were blocked in 5% non-fat milk/TBST and incubated with primary antibodies overnight at 4 °C, followed by HRP-conjugated secondary antibodies for 1 h at room temperature. Signals were detected by enhanced chemiluminescence (Bio-Rad) and visualized on X-ray film. Membranes were striped and re-probed with β-actin as loading control. Experiments were performed in triplicate.

### Cell viability assay

Cell viability was measured using an MTS assay (Abcam) per the manufacturer’s protocol. Cells were treated with indicated compounds for 48 h, incubated with 20 μL MTS reagent at 37 °C for 4 h, and absorbance was recorded at 490 nm using a microplate reader.

### Adenoviral constructs and production

INSM1, ASCL1, NEUROD1, NEUROG3, and MYCN cDNAs were cloned into *pShuttle-CMV* vectors (Ad5 backbone, E1/E3 deleted). Recombinant adenoviruses were generated using the AdEasy system (Agilent), with homologous recombination in E. coli BJ5183 and amplification in XL10-Gold cells. Viral particles were produced in AD-293 cells, purified by *CsCl* gradient centrifugation, and titrated using a hexon antibody (Thermo-Fisher Scientific, clone 3G0).

### INSM1 overexpression and knockdown in prostate cancer

INSM1 knockdown was achieved using adenoviral vectors encoding INSM1-specific *siRNA* (*5′-GGGAUCUGCUUAAAGUUUU-3′*) or scrambled control. Overexpression and knockdown efficiency were confirmed by Western blot of INSM1. All experiments were performed at least three times.

### Transcriptomic RAN-seq analysis of INSM1 overexpression and knockdown in prostate cancer

LNCaP cells were infected with adenovirus, *Ad-INSM1* for 48 hours to induce overexpression of INSM1. RNA sequencing was conducted using RNA extracted from LNCaP/*Ad-LacZ* and LNCaP/*Ad-INSM1* (each containing 2 biological replicates) at Novogene Corporation Inc (Sacramento, CA). The raw data were first proceeded using FASTP software to ensure data quality for further analysis. The raw data was submitted to public domain GEO (accession number GSE336536). The differential expression analysis was performed in R using the DESeq2 package. Gene with an adjust *p*-value less than 0.05 found by DESeq2 were assigned as significantly differential expression. The visualization heatmap was generated in R with heatmap package based on the results from DESeq2 analysis.

### Neuron-Related GO-BP Analysis

Differentially expressed genes from both NSM1 overexpression and INSM1 knockdown *RNA-seq* datasets were ranked according to DESeq2 Wald statistics and subjected to Gene Set Enrichment Analysis (GSEA) using the clusterProfiler package against Gene Ontology Biological Process (GO-BP) gene sets. Pathways with an adjusted *p-value (FDR) <0.05* were considered significant. Neuron-related pathways were identified using the keywords “nerv,” “neuro,” “axon,” “sensory,” “telencep,” and “endocrine.” Redundant GO terms were reduced by semantic similarity clustering using the Wang method in GOSemSim with a similarity cutoff of 0.6, retaining one representative term per cluster. Non-redundant neuronal GO-BP pathways were visualized according to normalized enrichment score (NES), gene set size, and adjusted p-value.

### Bioinformatic analysis

To examine the expression of INSM1 and NE markers across different prostate cancer states, *RNA-seq* data from patient cohorts (Beltran and SU2C) were obtained from cBioPortal (26, 55). Gene expression profiles from prostate cancer cell lines were derived from our previously published CTPC dataset (56) and were used for comparative analyses. *Single-cell RNA-seq* data were visualized using the Human Prostate Single Cell Atlas (HuPSA), which integrates datasets from multiple studies, including publicly available cohorts and data generated by our group (25). Epigenomic datasets were retrieved from the Gene Expression Omnibus (GEO), including *ATAC-seq* data in LNCaP and H660 cells (GSE139099) (57), INSM1 *ChIP-seq* data in pancreatic β-cells (GSE54046) (58), and ASCL1 *ChIP-seq* data in H660 cells (GSE183198) (35). All BigWig files were visualized using the Integrative Genomics Viewer (IGV) (59). Unless otherwise specified, datasets were aligned to the hg19 human genome assembly; INSM1 (mm10), H660, and ASCL1 datasets were aligned to hg38.

### Statistical analysis

Analyses were performed using GraphPad Prism v9.5.1. Data are presented as mean ± SD from ≥3 independent experiments, normalized to untreated controls where indicated. Two-group comparisons used unpaired two-tailed Student’s *t*-tests; multiple comparisons used one-way ANOVA with Tukey’s post hoc test. *p*<0.05 was considered significant (\**p*<0.05, \*\**p*<0.01, \*\*\**p*<0.001).

### CRediT authorship contribution statement

**Chiachen Chen:** Writing-original draft, methodology, formal analysis, and validation. **Siyuan Cheng:** Data curation, formal analysis, software, and methodology. **Lin Li:** Perform experiments. **Jeyaluxmy Sivalingam:** *RNA-seq* analysis, writing review, and editing. **Xin Gu:** Patient sample curation, perform histology experiment. **Yunshin Yeh:** Classified the Gleason Score, quantified the Allred score on human specimens. **Xiuping Yu:** Investigation, resources, supervision, writing review, and editing. **Michael S. Lan:** Conceptualization, funding acquisition, investigation, writing-review & editing, validation, and supervision.

## Acknowledgements

Authors acknowledge financial support from the Department of Genetics, LSUHSC and LSU Research Enhancement Program bridge grant funding to MSL. The FDA-approved oncology drugs set (AOD IX) was provided by the Drug Synthesis and Chemistry Branch, Developmental Therapeutics Program, Division of Cancer Treatment and Diagnosis, NCI (Bethesda, MD, USA).

## Data availability statement

The data are available from the authors on reasonable request. Data availability and deposition raw and processed *RNA-seq* data generated in this study have been deposited in the NCBI Gene Expression Omnibus (GEO) under accession number GSE336536. Raw sequencing reads (FASTQ format) were submitted via GEO’s Sequence Read Archive (SRA) pipeline using FTP/Aspera transfer. Each sample was annotated with metadata including organism, tissue source, library preparation method, sequencing platform, and experimental condition, following GEO’s metadata template requirements.

## Declaration of generative AI and AI-assisted technologies in the writing process

During the preparation of this work the authors used ChatGPT to improve readability. After using this tool, the authors reviewed and edited the content as needed and take full responsibility for the content of the publication.

## References

1. Yeh, Y., Guo, Q., Connelly, Z., Cheng, S., Yang, S., Prieto-Dominguez, N. et al. (2019) Wnt/Beta-Catenin Signaling and Prostate Cancer Therapy Resistance Adv Exp Med Biol 1210, 351–378

2. Beltran, H., Tomlins, S., Aparicio, A., Arora, V., Rickman, D., Ayala, G. et al. (2014) Aggressive variants of castration-resistant prostate cancer Clin Cancer Res 20, 2846–2850

3. Imamura, J., Ganguly, S., Muskara, A., Liao, R. S., Nguyen, J. K., Weight, C. et al. (2023) Lineage plasticity and treatment resistance in prostate cancer: the intersection of genetics, epigenetics, and evolution Front Endocrinol (Lausanne) 14, 1191311

4. Iwamoto, H., Nakagawa, R., Makino, T., Kadomoto, S., Yaegashi, H., Nohara, T. et al. (2022) Treatment Outcomes in Neuroendocrine Prostate Cancer Anticancer Res 42, 2167–2176

5. Brady, N. J., Bagadion, A. M., Singh, R., Conteduca, V., Van Emmenis, L., Arceci, E. et al. (2021) Temporal evolution of cellular heterogeneity during the progression to advanced AR-negative prostate cancer Nat Commun 12, 3372

6. Zhang, W., Liu, B., Wu, W., Li, L., Broom, B. M., Basourakos, S. P. et al. (2018) Targeting the MYCN-PARP-DNA Damage Response Pathway in Neuroendocrine Prostate Cancer Clin Cancer Res 24, 696–707

7. Storck, W. K., May, A. M., Westbrook, T. C., Duan, Z., Morrissey, C., Yates, J. A. et al. (2022) The Role of Epigenetic Change in Therapy-Induced Neuroendocrine Prostate Cancer Lineage Plasticity Front Endocrinol (Lausanne) 13, 926585

8. Jia, S., Wildner, H., and Birchmeier, C. (2015) Insm1 controls the differentiation of pulmonary neuroendocrine cells by repressing Hes1 Dev Biol 408, 90–98

9. Asrani, K., Torres, A. F., Woo, J., Vidotto, T., Tsai, H. K., Luo, J. et al. (2021) Reciprocal YAP1 loss and INSM1 expression in neuroendocrine prostate cancer J Pathol 255, 425–437

10. Breslin, M. B., Zhu, M., Notkins, A. L., and Lan, M. S. (2002) Neuroendocrine differentiation factor, IA-1, is a transcriptional repressor and contains a specific DNA-binding domain: identification of consensus IA-1 binding sequence. Nucleic Acids Res. 30, 1038–1045

11. Lan, M. S., and Breslin, M. B. (2009) Structure, expression, and biological function of INSM1 transcription factor in neuroendocrine differentiation. FASEB J. 23, 2024–2033

12. Mellitzer, G., Bonne, S., Luco, R. F., Van De Casteele, M., Lenne, N., Collombat, P., et al. (2006) IA-1 is Ngn3-dependent and essential for differentiation of the endocrine pancreas. EMBO J. 25, 1344–1352

13. Gierl, M. S., Karoulias, N., Wende, H., Strehle, M., and Birchmeier, C. (2006) The Zinc-finger factor Insm1 (IA-1) is essential for the development of pancreatic beta cells and intestinal endocrine cells. Genes Dev. 20, 2465–2478

14. Wildner, H., Gierl, M. S., Strehle, M., Pla, P., and Birchmeier, C. (2008) Insm1 (IA-1) is a crucial component of the transcriptional network that controls differentiation of the sympatho-adrenal lineage. Development 135, 473–481

15. Farkas, L. M., Haffner, C., Giger, T., Khaitovich, P., Nowick, K., Birchmeier, C. et al. (2008) Insulinoma-associated 1 has a panneurogenic role and promotes the generation and expansion of basal progenitors in the developing mouse neocortex. Neuron 60, 40–55

16. Chen, C., Notkins, A. L., and Lan, M . S. (2019) Insulinoma-Associated-1: From Neuroendocrine Tumor Marker to Cancer Therapeutics Mol Cancer Res 17, 1597–1604

17. Fujino, K., Motooka, Y., Hassan, W. A., Ali Abdalla, M. O., Sato, Y., Kudoh, S. et al. (2015) Insulinoma-Associated Protein 1 Is a Crucial Regulator of Neuroendocrine Differentiation in Lung Cancer *Am*.J.Pathol. 185, 3164–3177

18. Zhang, T., Liu, W. D., Saunee, N. A., Breslin, M. B., and Lan, M. S. (2009) Zinc-finger transcription factor INSM1 interrupts cyclin D1 and CDK4 binding and induces cell cycle arrest. J.Biol.Chem. 284, 5574–5581

19. Rosenbaum, J. N., Guo, Z., Baus, R. M., Werner, H., Rehrauer, W. M., and Lloyd, R. V. (2015) INSM1: A Novel Immunohistochemical and Molecular Marker for Neuroendocrine and Neuroepithelial Neoplasms *Am*.J.Clin.Pathol. 144, 579–591

20. Chen, C., Breslin, M. B., and Lan, M. S. (2018) Sonic hedgehog signaling pathway promotes INSM1 transcription factor in neuroendocrine lung cancer Cell Signal 46, 83–91

21. Chen, C., Breslin, M. B., Guidry, J. J., and Lan, M. S. (2019) 5’-Iodotubercidin represses insulinoma-associated-1 expression, decreases cAMP levels, and suppresses human neuroblastoma cell growth J Biol Chem 294, 5456–5465

22. Chen, C., Wu, J., Hicks, C., and Lan, M. S. (2023) Repurposing a plant alkaloid homoharringtonine targets insulinoma associated-1 in N-Myc-activated neuroblastoma Cell Signal 109, 110753

23. Chen, C., and Lan, M. S. (2020) A promoter-driven assay for INSM1-associated signaling pathway in neuroblastoma Cellular signalling 76, 109785

24. Cheng, S., and Yu, X. (2019) Bioinformatics analyses of publicly available NEPCa datasets Am J Clin Exp Urol 7, 327–340

25. Cheng, S., Li, L., Yeh, Y., Shi, Y., Franco, O., Corey, E., et al. (2024) Unveiling novel double-negative prostate cancer subtypes through single-cell RNA sequencing analysis NPJ Precis Oncol 8, 171

26. Gao, J., Aksoy, B. A., Dogrusoz, U., Dresdner, G., Gross, B., Sumer, S. O., et al. (2013) Integrative analysis of complex cancer genomics and clinical profiles using the cBioPortal Sci Signal 6, pl1

27. Lee, J. K., Phillips, J. W., Smith, B. A., Park, J. W., Stoyanova, T., McCaffrey, E. F. et al. (2016) N-Myc Drives Neuroendocrine Prostate Cancer Initiated from Human Prostate Epithelial Cells Cancer Cell 29, 536–547

28. Ji, Y., Zhang, W., Shen, K., Su, R., Liu, X., Ma, Z., et al. (2023) The ELAVL3/MYCN positive feedback loop provides a therapeutic target for neuroendocrine prostate cancer Nat Commun 14, 7794

29. Zertal-Zidani, S., Bounacer, A., and Scharfmann, R. (2007) Regulation of pancreatic endocrine cell differentiation by sulphated proteoglycans Diabetologia 50, 585–595

30. Bizzaro, C. L., Bach, C. A., Santos, R. A., Verrillo, C. E., Naranjo, N. M., Chaudhari, I. et al. (2025) Exploring STEAP1 Expression in Prostate Cancer Cells in Response to Androgen Deprivation and in Small Extracellular Vesicles Mol Cancer Res 23, 542–552

31. Li, L., Xu, Q., and Tang, C. (2023) RGS proteins and their roles in cancer: friend or foe? Cancer Cell Int 23, 81

32. Sivalingam, J., Cho, K. H., Shi, Y., Barron, P., Lan, M., Franco, O., et al. (2025) A MYC Family Switch: L-MYC Drives and Maintains Neuroendocrine Lineage Programs in Prostate Cancer bioRxiv 10.64898/2025.12.25.696507

33. Cejas, P., Xie, Y., Font-Tello, A., Lim, K., Syamala, S., Qiu, X., et al. (2021) Subtype heterogeneity and epigenetic convergence in neuroendocrine prostate cancer Nat Commun 12, 5775

34. Rudin, C. M., Poirier, J. T., Byers, L. A., Dive, C., Dowlati, A., George, J. et al. (2019) Molecular subtypes of small cell lung cancer: a synthesis of human and mouse model data Nat Rev Cancer 19, 289–297

35. Nouruzi, S., Ganguli, D., Tabrizian, N., Kobelev, M., Sivak, O., Namekawa, T. et al. (2022) ASCL1 activates neuronal stem cell-like lineage programming through remodeling of the chromatin landscape in prostate cancer Nat Commun 13, 2282

36. Lan, M. S., and Chen, C. (2023) Small Molecules Targeting INSM1 for the Treatment of High-Risk Neuroblastoma Biology (Basel) 12,

37. Alvandi, F., Kwitkowski, V. E., Ko, C. W., Rothmann, M. D., Ricci, S., Saber, H. et al. (2014) U.S. Food and Drug Administration approval summary: omacetaxine mepesuccinate as treatment for chronic myeloid leukemia Oncologist 19, 94–99

38. Levy, V., Zohar, S., Bardin, C., Vekhoff, A., Chaoui, D., Rio, B. et al. (2006) A phase I dose-finding and pharmacokinetic study of subcutaneous semisynthetic homoharringtonine (ssHHT) in patients with advanced acute myeloid leukaemia Br J Cancer 95, 253–259

39. Liu, D., Xing, J., Xiong, F., Yang, F., and Gu, N. (2017) Preparation and in vivo safety evaluations of antileukemic homoharringtonine-loaded PEGylated liposomes Drug Dev Ind Pharm 43, 652–660

40. Davies, A. H., Beltran, H., and Zoubeidi, A. (2018) Cellular plasticity and the neuroendocrine phenotype in prostate cancer Nat Rev Urol 15, 271–286

41. Conteduca, V., Oromendia, C., Eng, K. W., Bareja, R., Sigouros, M., Molina, A. et al. (2019) Clinical features of neuroendocrine prostate cancer Eur J Cancer 121, 7–18

42. Wang, Y., Wang, Y., Ci, X., Choi, S. Y. C., Crea, F., Lin, D. et al. (2021) Molecular events in neuroendocrine prostate cancer development Nat Rev Urol 18, 581–596

43. Wang, H. T., Yao, Y. H., Li, B. G., Tang, Y., Chang, J. W., and Zhang, J. (2014) Neuroendocrine Prostate Cancer (NEPC) progressing from conventional prostatic adenocarcinoma: factors associated with time to development of NEPC and survival from NEPC diagnosis-a systematic review and pooled analysis J Clin Oncol 32, 3383–3390

44. Cacciatore, A., Albino, D., Catapano, C. V., and Carbone, G. M. (2023) Preclinical Models of Neuroendocrine Prostate Cancer *Curr Protoc* **3**, e742

45. Epstein, J. I., Amin, M. B., Beltran, H., Lotan, T. L., Mosquera, J. M., Reuter, V. E. et al. (2014) Proposed morphologic classification of prostate cancer with neuroendocrine differentiation Am J Surg Pathol 38, 756–767

46. Zhang, T., Wang, H. W., Saunee, N. A., Breslin, M. B., and Lan, M. S. (2010) Insulinoma-associated antigen-1 zinc-finger transcription factor promoters pancreatic duct cell trans-differentiation. Endocrinology 151, 2030–2039

47. Zhang, T., Saunee, N. A., Breslin, M. B., Song, K., and Lan, M. S. (2012) Functional role of an islet transcription factor, INSM1/IA-1, on pancreatic acinar cell trans-differentiation. J.Cell.Physiol. 227, 2470–2479

48. Zhu, M., Breslin, M. B., and Lan, M. S. (2002) Expression of a novel zinc-finger cDNA, IA-1, is associated with rat AR42J cells differentiation into insulin-positive cells. Pancreas 24, 139–145

49. Chen, C., Cheng, S., Yu, X., Lee, Y., and Lan, M. S. (2025) Regulation of INSM1 Gene Expression and Neuroendocrine Differentiation in High-Risk Neuroblastoma Biology (Basel) 15, 22

50. Shimomura, T., Murakami, M., Kasai, K., Imai, Y., Inaba, Y., Urabe, F. et al. (2026) Assessment of Circulating Neuroendocrine Tumor Markers at Prostate Cancer Diagnosis: An Investigation of Prostate Cancer With Neuroendocrine Features Prostate 86, 1035–1042

51. Zitella, A., Berruti, A., Destefanis, P., Mengozzi, G., Torta, M., Ceruti, C. et al. (2007) Comparison between two commercially available chromogranin A assays in detecting neuroendocrine differentiation in prostate cancer and benign prostate hyperplasia Clin Chim Acta 377, 103–107

52. Yinjun, L., Jie, J., Weilai, X., and Xiangming, T. (2004) Homoharringtonine mediates myeloid cell apoptosis via upregulation of pro-apoptotic bax and inducing caspase-3-mediated cleavage of poly(ADP-ribose) polymerase (PARP) American journal of hematology 76, 199–204

53. Zhang, T., Shen, S., Zhu, Z., Lu, S., Yin, X., Zheng, J. et al. (2016) Homoharringtonine binds to and increases myosin-9 in myeloid leukaemia British journal of pharmacology 173, 212–221

54. Cheng, S., Prieto-Dominguez, N., Yang, S., Connelly, Z. M., StPierre, S., Rushing, B. et al. (2020) The expression of YAP1 is increased in high-grade prostatic adenocarcinoma but is reduced in neuroendocrine prostate cancer Prostate Cancer Prostatic Dis 23, 661–669

55. Cerami, E., Gao, J., Dogrusoz, U., Gross, B. E., Sumer, S. O., Aksoy, B. A. et al. (2012) The cBio cancer genomics portal: an open platform for exploring multidimensional cancer genomics data Cancer Discov 2, 401–404

56. Cheng, S., and Yu, X. (2023) CTPC, a combined transcriptome data set of human prostate cancer cell lines Prostate 83, 158–161

57. Grubert, F., Srivas, R., Spacek, D. V., Kasowski, M., Ruiz-Velasco, M., Sinnott-Armstrong, N. et al. (2020) Landscape of cohesin-mediated chromatin loops in the human genome Nature 583, 737–743

58. Jia, S., Ivanov, A., Blasevic, D., Muller, T., Purfurst, B., Sun, W. et al. (2015) Insm1 cooperates with Neurod1 and Foxa2 to maintain mature pancreatic beta-cell function EMBO J 34, 1417–1433

59. Robinson, J. T., Thorvaldsdottir, H., Winckler, W., Guttman, M., Lander, E. S., Getz, G. et al. (2011) Integrative genomics viewer Nat Biotechnol 29, 24–26

